# Sample-Index Misassignment Impacts Tumor Exome Sequencing

**DOI:** 10.1101/182659

**Authors:** Daniel Vodák, Susanne Lorenz, Sigve Nakken, Lars Birger Aasheim, Harald Holte, Baoyan Bai, Ola Myklebost, Leonardo A. Meza-Zepeda, Eivind Hovig

## Abstract

Sample pooling enabled by dedicated indexes is a common and cost-effective strategy used in high-throughput DNA sequencing. Index misassignment leading to cross-sample contamination has however been described as a general problem of sequencing instruments which utilize exclusion amplification. Using real-life data from multiple tumor sequencing projects, we demonstrate that co-multiplexed samples can induce artifactual calls closely resembling high-quality somatic variant calls, and argue that dual indexing is the most reliable countermeasure.

---

Identification of somatic variants by next-generation sequencing has become an important technique in cancer research by pinpointing the genomic causes of tumor phenotypes. An increasing number of examples have further shown that genomic aberrations have prognostic value and can inform rational clinical deployment of targeted cancer drugs (Garraway, 2013; Griffith *et al.*, 2017). As of now, next-generation sequencing technologies enable affordable assaying of variation in the entire tumor genome within days, but despite the availability of specialized software tools, somatic variant calling continues to pose challenges. Notably, the inherent complexity of tumors, as exemplified by aneuploidy, tumor heterogeneity, and sample impurity, often leads to important somatic variants only being detectable in low allelic fractions (AF) (Cibulskis *et al.*, 2013; Ryu *et al.*, 2016; Lee *et al.*, 2016; Cheng *et al.*, 2015; Zehir *et al.*, 2017). AFs can be further affected due to technical reasons, such as the inability to accurately represent allelic ratios at genomic loci with low coverage (Carter *et al.*, 2012). As a consequence of the necessarily high required sensitivity, somatic variant calling is susceptible to random noise, systematic artefacts, and sample contamination (Alioto **et al.**, 2015; Chen *et al.*, 2017). When unnoticed, false positive variant calls can contribute to high costs incurred by follow-up analyses and experiments, and in the worst case support inadequate therapeutic strategies.

As originally described in the context of RNA-seq applications (Sinha *et al.*, 2017), the sequencing technology itself can be a source of sample cross-contamination when material from multiple samples is being “barcoded” by dedicated indexes and pooled together before sequencing. Pronounced on Illumina instruments utilizing exclusion amplification (ExAmp) and patterned flow cells (i.e., HiSeq 3000/4000/X Ten and NovaSeq), sample index misassignment leads to transfer of individual sequencing reads between samples multiplexed into a shared sequencing pool. Our analysis of real-life data shows that index misassignment is a source of false positive somatic variant calls in a form of true variation obtained from co-multiplexed samples.

We investigated tumor-normal sample pairs of three different tumor types: diffuse large B-cell lymphoma (DLBCL), follicular lymphoma (FL), and sarcoma (SARC). All samples were collected from cancer patients in Norway. Multiple Illumina instruments were used for deep exome sequencing of the selected samples (median coverage: 315X for tumor samples, 146X for controls): three HiSeq 2000/2500 instruments utilizing bridge amplification (generating 42 DLBCL and 31 SARC tumor samples) and a HiSeq 4000 employing ExAmp (generating 81 FL and 77 SARC tumor samples). All samples were subject to standardized library preparation, sequencing, and bioinformatic analysis, as adopted by the Norwegian Cancer Genomics Consortium (NCGC, http://cancergenomics.no) (Supplementary methods). In our assessment of index misassignment, we have not addressed its causes, but focused on consequences for calling of somatic single nucleotide variants (sSNV). Only sSNVs agreed upon by two independent variant callers (MuTect and Strelka) were considered (Cibulskis **et al.**, 2013; Saunders *et al.*, 2012). Sample-wise contamination estimates were generated by Conpair (Bergmann *et al.*, 2016), which assesses the allelic composition on several thousand genomic markers. Our tests on simulated data suggest that Conpair provides consistent quantifications, despite apparent progressive underestimation dependent on the number of contributing contamination sources (Supplementary Table 1, Supplementary methods). We attribute this underestimation to decreasing germline variant profile concordance in sample pools of increasing size (Supplementary Fig. 1).

During the analysis of hundreds of sequenced tumor-normal matched pairs, we noted that data generated on the ExAmp instrument showed significantly higher sample-wise contamination levels in comparison to data from bridge amplification instruments (median per-sample contamination estimate values: 0.839% vs. 0.187%, p-value < 0.001, Supplementary Fig. 2). To specifically assess the role of multiplexing on the ExAmp instrument, we sequenced a selected set of 16 sample libraries both in pools and individually (i.e., one sample library per flow cell lane), observing significantly increased contamination rates in datasets from the multiplexed libraries (median per-sample contamination estimate values: 0.644% vs. 0.0465%, p-value < 0.001, Supplementary Table 2). Removal of free adapters/primers prior to sequencing has been identified as a key measure for mitigating the cross-contamination rates (Illumina, 2017). For the 16 libraries included in our testing, performing a gel-based library purification step in combination with bead purification did not provide improvements over bead purification alone (median per-sample contamination estimate values: 0.671% vs. 0.6115%, p-value = 0.159). In accordance with previous reports by Illumina and the study by Sinha *et al.*, we concluded that ExAmp chemistry is the cause of sample contamination. The dependency on sample multiplexing indicated that co-multiplexed samples serve as the contaminants.

To estimate the impact of ExAmp-associated contamination on calling of sSNVs, we calculated the read fraction support of each variant in its respective sample pool of origin. By considering only reads assigned to samples of other individuals, we were able to focus strictly on inter-individual contamination. Hence, we denote the set of reads considered for a given variant during our calculations as an “individual-based pool complement” (IBPC) (Supplementary Fig. 3). Two distinct classes of variants can be identified when variant AFs are plotted against variant read support in respective IBPCs: 1) apparently true somatic variants (ATSVs), consisting of variants not present in the Norwegian population (i.e. absent from NCGC’s cohort of normal samples) and lacking support in the IBPC (present in <1% of variant site IBPC reads); and 2) suspected contaminant variants (SCVs), consisting of common Norwegian germline variants (>= 5% allele frequency in NCGC’s cohort of normal samples) while also having a considerable support in IBPC (present in >=20% of variant site IBPC reads) (Supplementary Fig. 4). We classify the remaining variants as ambiguous. In comparison to bridge amplification, ExAmp leads to higher occurrence of sSNVs that coincide with germline variation common in the Norwegian population (Fig. 1). IBPC read fraction support for these variants is significantly higher in the ExAmp datasets (median values: 0.508 vs. 0.125, p-value < 0.001), showing a much better correspondence to the suspected contamination source. Samples sequenced on the ExAmp instrument show significantly higher SCV counts than samples sequenced on bridge amplification instruments (per-sample median counts: 4 vs. 0, p-value < 0.001) and significant correspondence between the number of SCVs and the estimated contamination (p-values < 0.001, Fig. 2).

**Figure 1.**
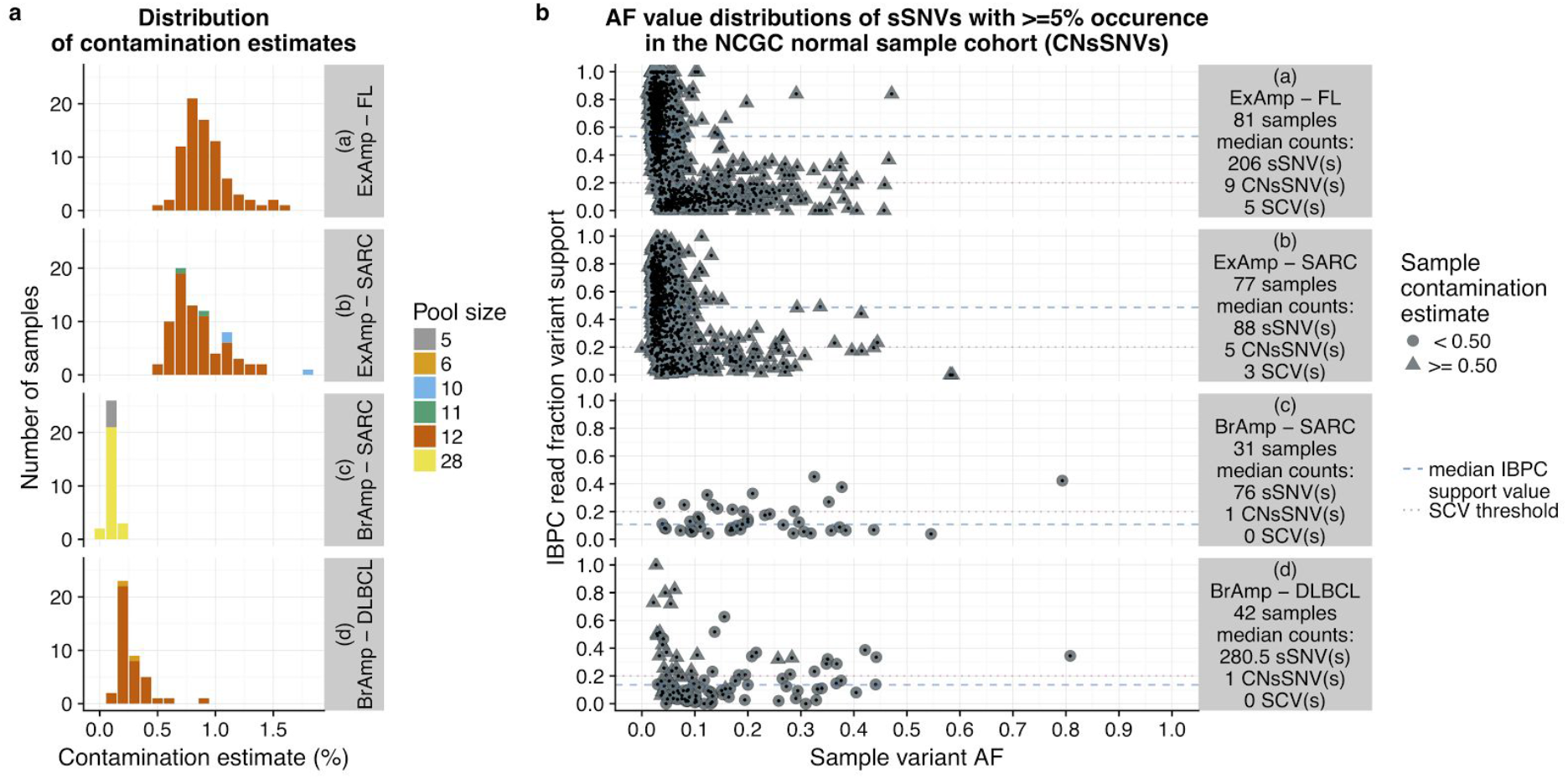
The relationships between amplification chemistry, sample contamination estimates, and IBPC read fraction variant support. Values are plotted separately for each combination of tumor type and amplification chemistry represented in the analysis. BrAmp: bridge amplification.

**Figure 2.**
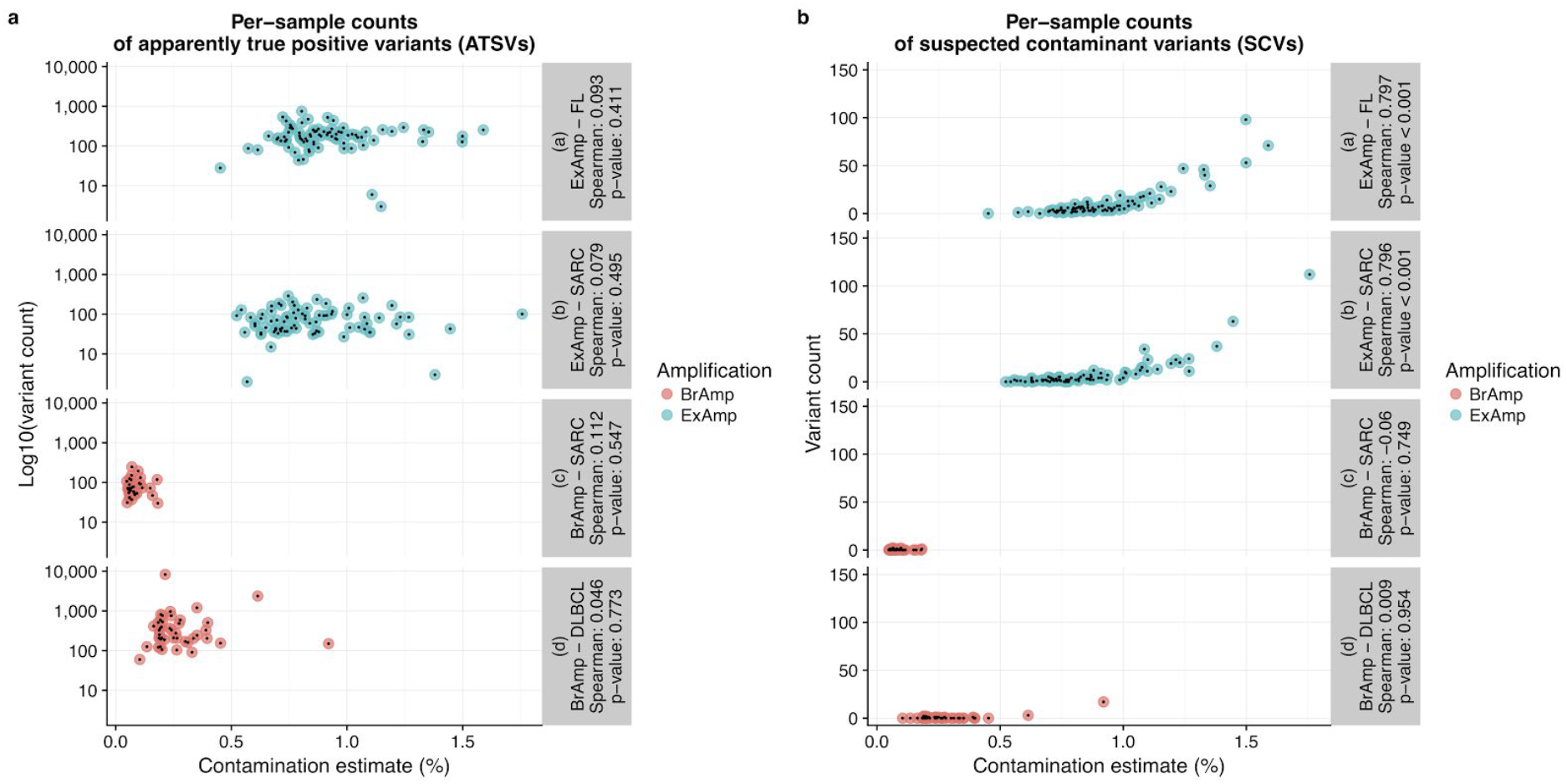
Apparently true somatic variants (A) and suspected contaminant variants (B). Per-sample variant counts are plotted separately for each combination of tumor type and amplification chemistry represented in the analysis.

We have focused on exploring the general link between cross-sample contamination levels and detection of SCVs with germline origin, but additional factors could be considered. Cross-sample contamination by recurrent somatic variants would represent a class of potential false positives that would be more difficult to recognize, and may have stronger clinical implications, considering that some actionable variants occur at high frequency across tumor samples. Concern for such cases of contamination gains relevance when co-multiplexing samples with identical somatic variants (e.g., pooling serial patient biopsies or screening multiple samples likely to harbor hotspot mutations), particularly in sensitive analyses of high purity samples. When co-multiplexing tumor and control samples, false negative variant rates might be increased due to somatic variation contaminating the controls. Moreover, in copy number loss regions of high purity tumor samples, we expect both the number and the average AF of SCVs to be increased due to the lack of non-contaminant reads.

Several countermeasures have been suggested for the prevention of index misassignment, with sample storage conditions being shown by Illumina to influence the problem’s severity. Sequencing of a single sample per lane proved to be an effective solution, which however might be too costly for many applications. Rigorous gel-and bead-based library purification have been strongly recommended as means of index misassignment mitigation in case of sample multiplexing, but appear to be insufficient even in combination. Dual indexing has been suggested for circumventing the problem by using sample-specific index pairs rather than individual indexes, enabling recognition of reads with unexpected index combinations caused by index misassignment. Dual indexes are becoming a part of best practices for multiplexed libraries sequenced with ExAmp chemistry, but their current availability may vary depending on the library preparation protocol. The possible solutions therefore need to be assessed in the context of each particular project. For data that have already been generated, strict filtering based on IBPC values could provide a quick solution. A more sophisticated filtering approach might be desirable in order to limit false negative rates, particularly among variants with low AFs, a category that harbors variants of prognostic and therapeutic value in multiple cancer types (Landau *et al.*, 2013; McGranahan *et al.*, 2015; Schmitt *et al.*, 2016).

We believe our findings to be of relevance to other cancer sequencing projects that utilize the ExAmp chemistry, even though the impacts of index misassignment on somatic variant calling may vary depending on the combination of employed sequencing instruments, library preparation protocols, and bioinformatic analyses. We note that besides single nucleotide variants, other types of somatic variation (e.g. insertions and deletions) are also likely to be affected.

## Acknowledgements

We are grateful for the sequencing services provided by the Genomics Core Facility, South East Health Region/Oslo University Hospital, and for technical assistance provided by Jinchang Sun. We thank Illumina, Inc. for providing sequencing reagents and flow cells necessary to conduct the experiment for evaluating the role of multiplexing in ExAmp-associated index misassignment. This work has been supported by the Research Council of Norway (RCN grants 218241, 221580 and 230817/F20)

## Author contributions

DV and SL designed data analysis. DV, SN and LBA conducted data analysis. DV, SL, LAM-Z and EH designed the sequencing experiment. DV and SN drafted the manuscript. SL, LBA, OM, HH, BB, LAM-Z and EH revised the manuscript. OM, HH and BB acquired sample data. EH and LAM-Z supervised the project. All authors read and approved the final manuscript.

---

**Supplementary Figure 1.**
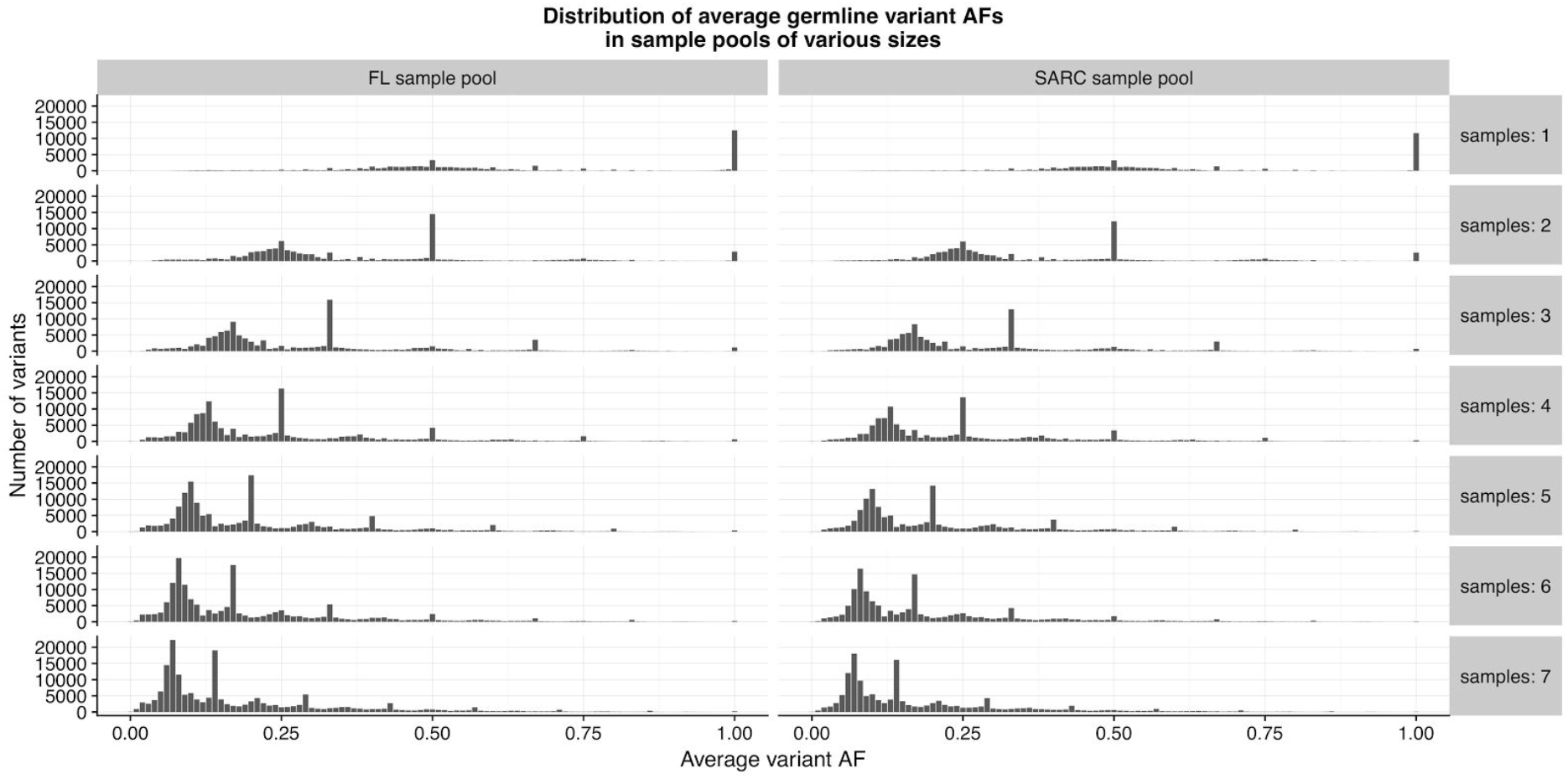
Distribution of average germline single nucleotide variant AFs in sample pools of various sizes. Plotted are only potential contamination variants (i.e., variants not present in an additional sample representing the target of contamination) from each given pool. As the number of samples (i.e., potential contamination sources) in a given pool increases, the average AF decreases for most of the present germline variants, diluting the contamination potential.

**Supplementary Figure 2.**
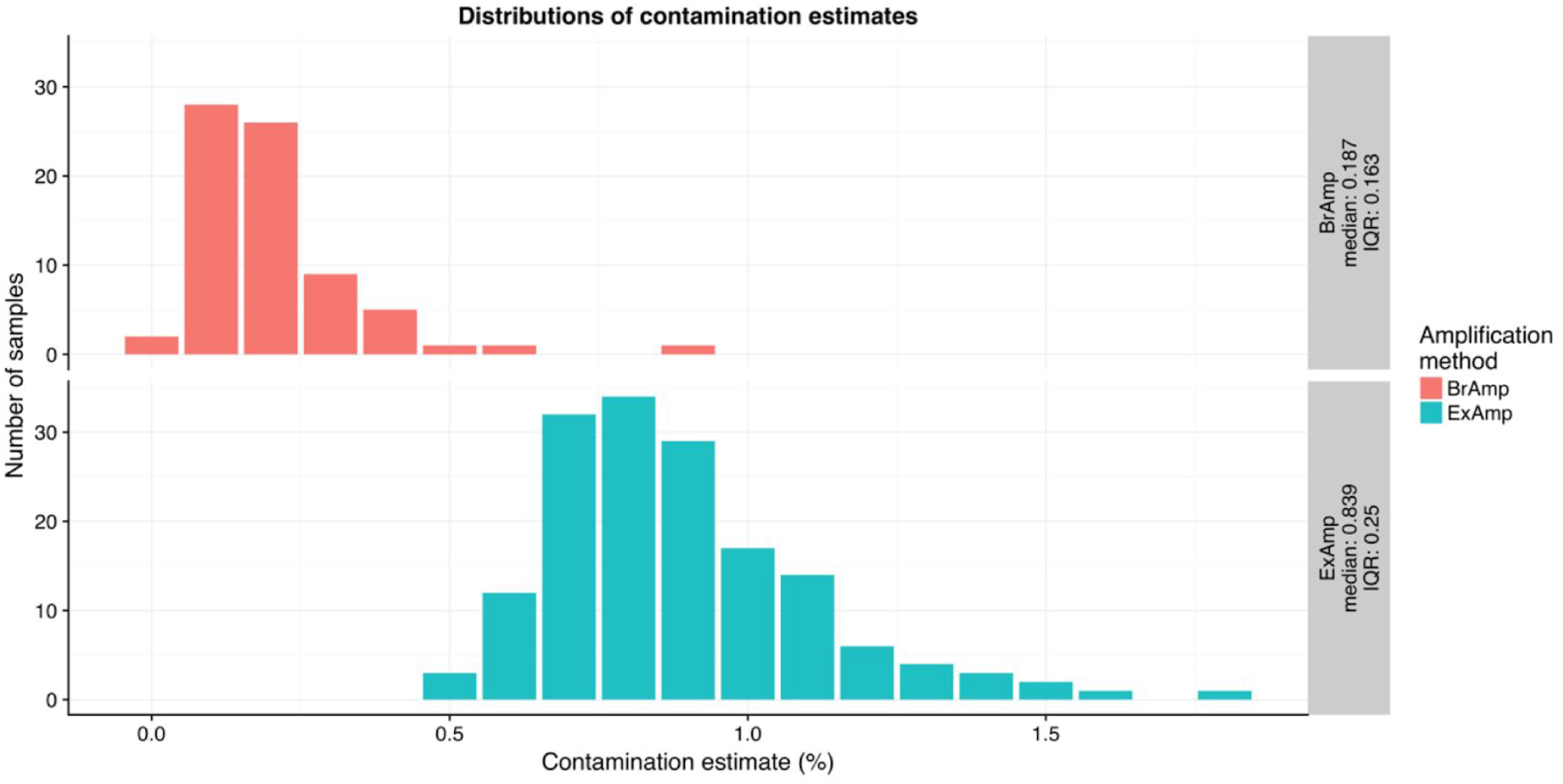
Distributions of tumor sample contamination estimates generated by Conpair. Separate distributions are plotted for samples sequenced with bridge amplification instruments (top) and samples sequenced with an exclusion amplification instrument (bottom).

**Supplementary Figure 3.**
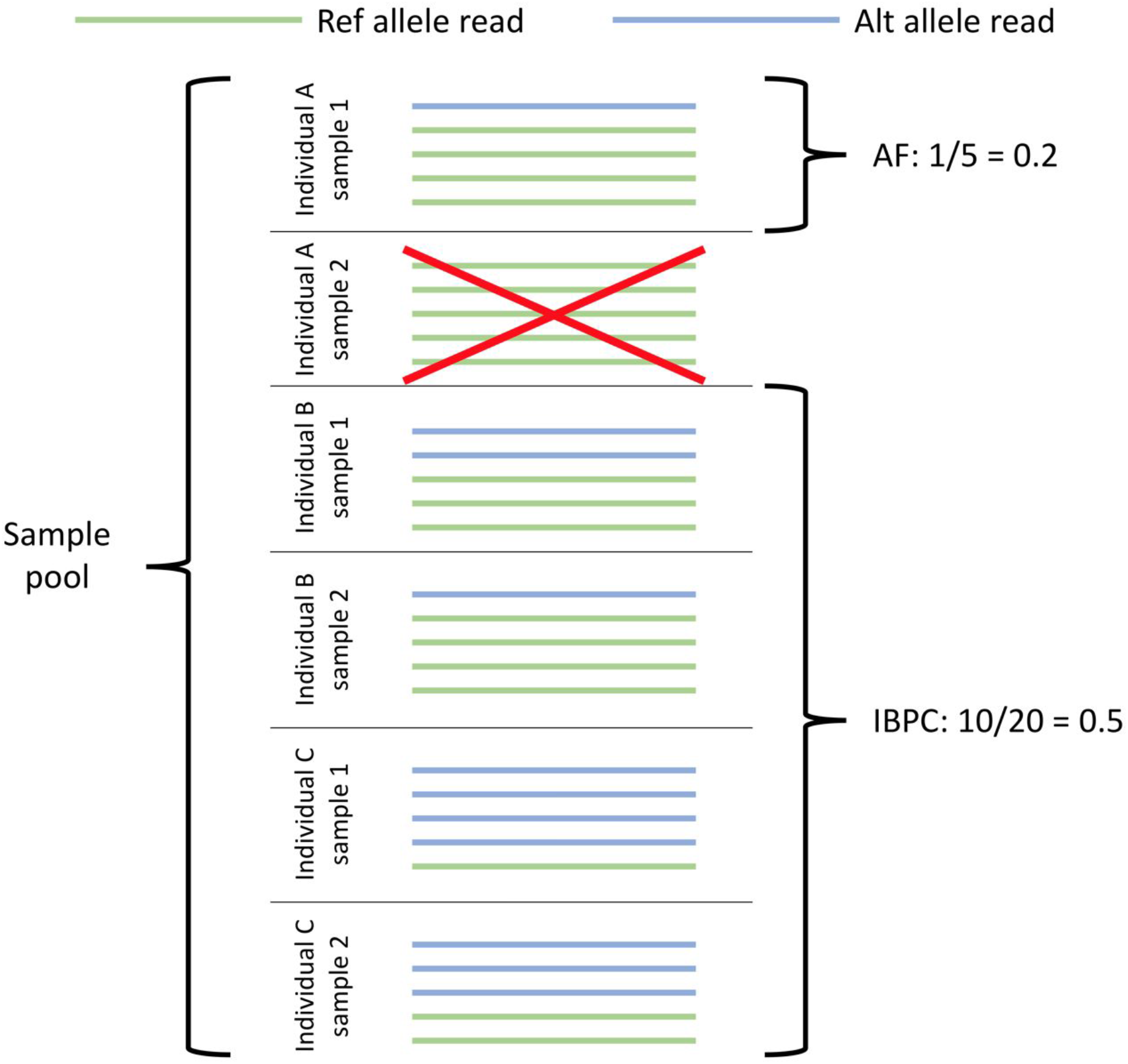
Calculation of IBPC read fraction variant support values. When calculating IBPC values for variants called in a sample of a given individual (e.g., sample 1 of individual A), only reads assigned to samples of other individuals are considered.

**Supplementary Figure 4.**
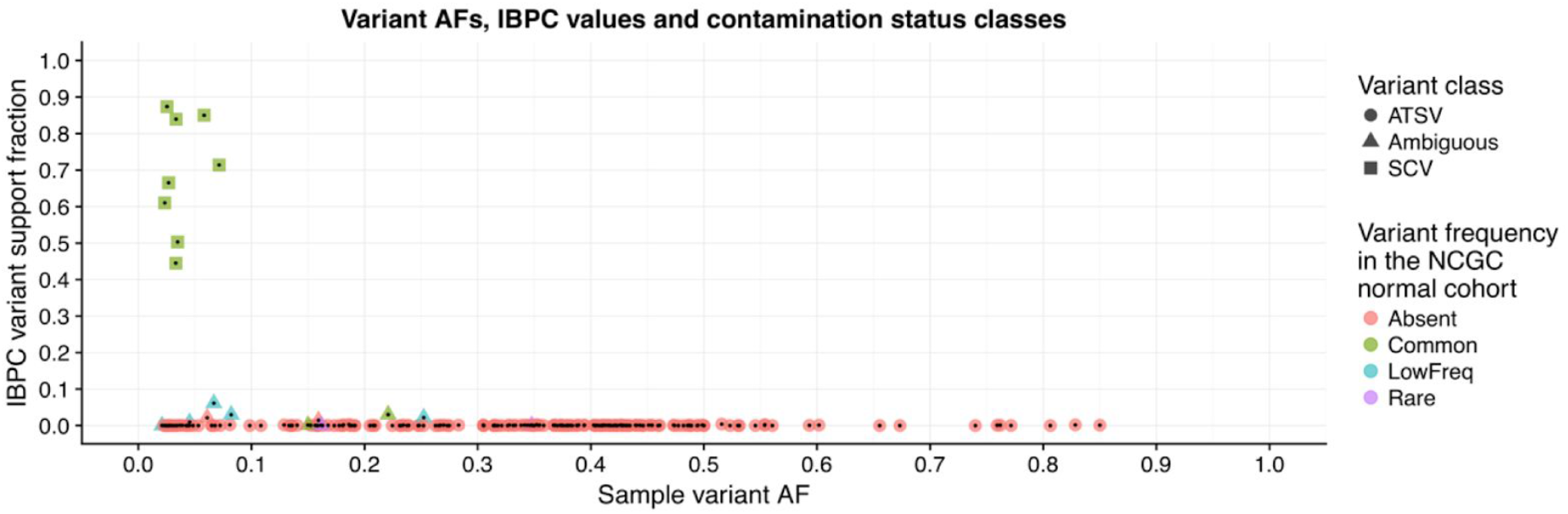
Classification of sSNVs according to their contamination status (example for a selected tumor sample). ATSVs are defined by IBPC variant support fraction < 0.01 and absence in the NCGC cohort of normals. SCVs appear in the NCGC cohort of normals with >= 5% frequency and have IBPC variant support fraction >= 0.2. Ambiguous cases (all remaining variants) are typically also present; they are composed of recurrent somatic variants, true somatic variants appearing at germline variant locations, and false positive calls potentially unrelated to index misassignment.

**Supplementary Table 1.** Accuracy of contamination estimates generated by Conpair on simulated data. Reads from multiple libraries were mixed to simulate contamination by 1, 2, 4 or 7 samples from other individuals, with the total contamination amounting to ~8% in each case. The right-most column shows how much larger the true contamination content is compared to the corresponding estimate by Conpair.

| Contaminated sample | Number of<br>contaminating<br>samples | Conpair's<br>estimate of<br>contamination<br>(%) | True<br>contamination<br>content (%) | True contamination<br>content as a fraction of<br>Conpair's estimate of<br>contamination |
| --- | --- | --- | --- | --- |
| B_ExAmp_SARC_0013_tumor | 1 | 8.043 | 7.999 | 0.99 |
| B_ExAmp_SARC_0016_tumor | 1 | 8.217 | 8.001 | 0.97 |
| B_ExAmp_SARC_0021_tumor | 1 | 8.663 | 7.993 | 0.92 |
| B_ExAmp_SARC_0030_tumor | 1 | 8.277 | 8.01 | 0.97 |
| B_ExAmp_SARC_0051_tumor | 1 | 7.812 | 7.995 | 1.02 |
| B_ExAmp_SARC_0057_tumor | 1 | 8.081 | 7.998 | 0.99 |
| B_ExAmp_SARC_0063_tumor | 1 | 8.412 | 8.009 | 0.95 |
| B_ExAmp_SARC_0013_tumor | 2 | 6.196 | 7.999 | 1.29 |
| B_ExAmp_SARC_0016_tumor | 2 | 6.371 | 7.992 | 1.25 |
| B_ExAmp_SARC_0021_tumor | 2 | 6.504 | 7.981 | 1.23 |
| B_ExAmp_SARC_0030_tumor | 2 | 6.688 | 7.98 | 1.19 |
| B_ExAmp_SARC_0051_tumor | 2 | 5.988 | 8.001 | 1.34 |
| B_ExAmp_SARC_0057_tumor | 2 | 6.293 | 7.988 | 1.27 |
| B_ExAmp_SARC_0063_tumor | 2 | 6.286 | 8.003 | 1.27 |
| B_ExAmp_SARC_0013_tumor | 4 | 5.676 | 7.989 | 1.41 |
| B_ExAmp_SARC_0016_tumor | 4 | 5.707 | 7.995 | 1.4 |
| B_ExAmp_SARC_0021_tumor | 4 | 5.743 | 8.014 | 1.4 |
| B_ExAmp_SARC_0030_tumor | 4 | 6.244 | 8.001 | 1.28 |
| B_ExAmp_SARC_0051_tumor | 4 | 5.414 | 7.986 | 1.48 |
| B_ExAmp_SARC_0057_tumor | 4 | 5.607 | 7.967 | 1.42 |
| B_ExAmp_SARC_0063_tumor | 4 | 5.532 | 7.985 | 1.44 |
| B_ExAmp_SARC_0013_tumor | 7 | 5.54 | 7.996 | 1.44 |
| B_ExAmp_SARC_0016_tumor | 7 | 5.495 | 8.001 | 1.46 |
| B_ExAmp_SARC_0021_tumor | 7 | 5.605 | 7.996 | 1.43 |
| B_ExAmp_SARC_0030_tumor | 7 | 6.091 | 8.004 | 1.31 |
| B_ExAmp_SARC_0051_tumor | 7 | 5.27 | 8.014 | 1.52 |
| B_ExAmp_SARC_0057_tumor | 7 | 5.411 | 8.007 | 1.48 |
| B_ExAmp_SARC_0063_tumor | 7 | 5.412 | 8.016 | 1.48 |

**Supplementary Table 2.**
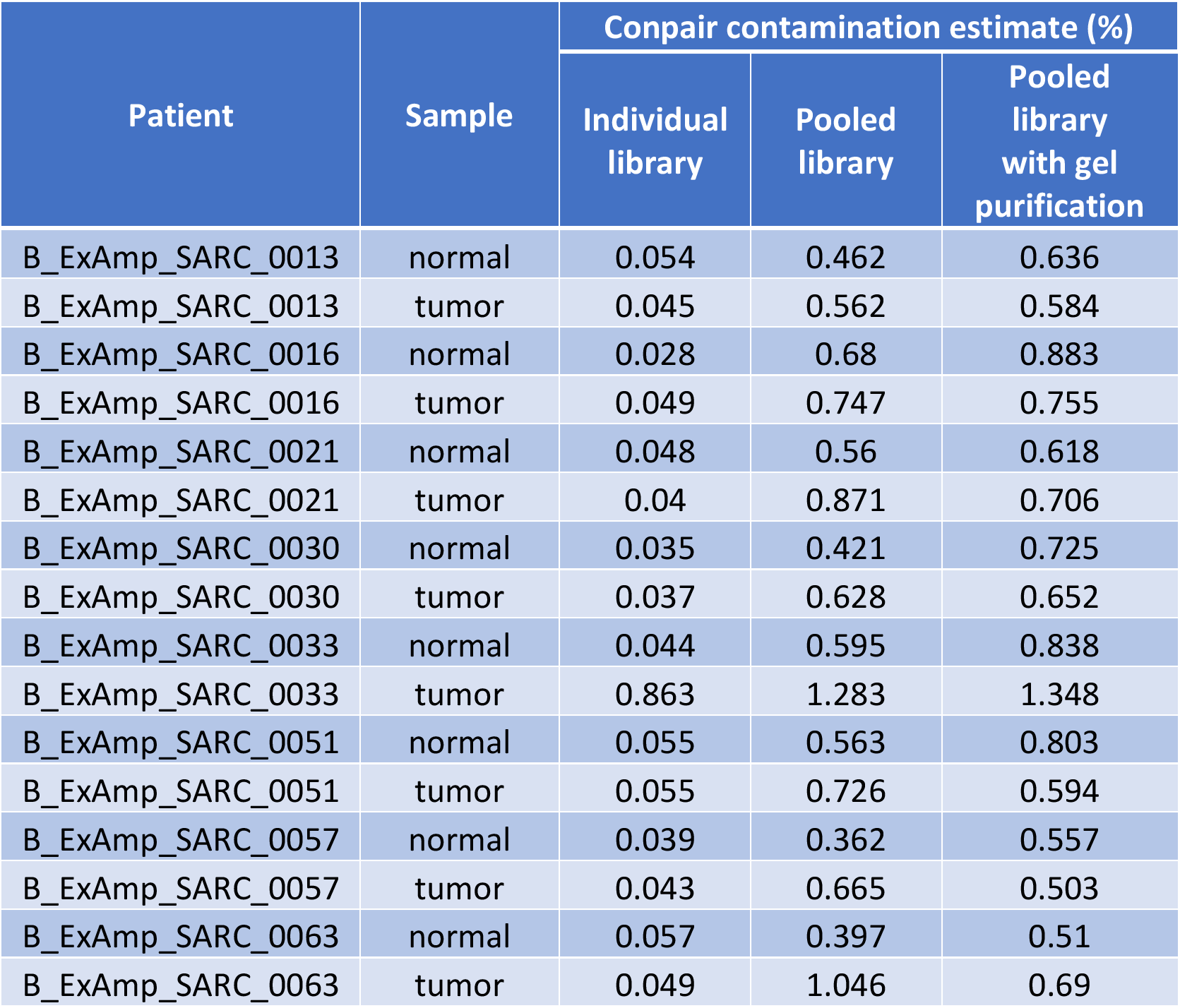
Conpair contamination estimates for sample libraries sequenced one-per-lane *(Individual library)* and in eight-sample pools *(Pooled library* and *Pooled library with gel purification)* on an ExAmp instrument.

| Patient | Sample | Conpair contamination estimate (%) |  |  |
| --- | --- | --- | --- | --- |
|  |  | Individual library | Pooled library | Pooled library with gel purification |
| B_ExAmp_SARC_0013 | normal | 0.054 | 0.462 | 0.636 |
| B_ExAmp_SARC_0013 | tumor | 0.045 | 0.562 | 0.584 |
| B_ExAmp_SARC_0016 | normal | 0.028 | 0.68 | 0.883 |
| B_ExAmp_SARC_0016 | tumor | 0.049 | 0.747 | 0.755 |
| B_ExAmp_SARC_0021 | normal | 0.048 | 0.56 | 0.618 |
| B_ExAmp_SARC_0021 | tumor | 0.04 | 0.871 | 0.706 |
| B_ExAmp_SARC_0030 | normal | 0.035 | 0.421 | 0.725 |
| B_ExAmp_SARC_0030 | tumor | 0.037 | 0.628 | 0.652 |
| B_ExAmp_SARC_0033 | normal | 0.044 | 0.595 | 0.838 |
| B_ExAmp_SARC_0033 | tumor | 0.863 | 1.283 | 1.348 |
| B_ExAmp_SARC_0051 | normal | 0.055 | 0.563 | 0.803 |
| B_ExAmp_SARC_0051 | tumor | 0.055 | 0.726 | 0.594 |
| B_ExAmp_SARC_0057 | normal | 0.039 | 0.362 | 0.557 |
| B_ExAmp_SARC_0057 | tumor | 0.043 | 0.665 | 0.503 |
| B_ExAmp_SARC_0063 | normal | 0.057 | 0.397 | 0.51 |
| B_ExAmp_SARC_0063 | tumor | 0.049 | 1.046 | 0.69 |

## Reference

Alioto, T.S. et al. (2015) A comprehensive assessment of somatic mutation detection in cancer using whole-genome sequencing. Nat. Commun., 6, 10001.

Bergmann, E.A. et al. (2016) Conpair: concordance and contamination estimator for matched tumor-normal pairs. Bioinformatics, 32, 3196-3198.

Carter, S.L. et al. (2012) Absolute quantification of somatic DNA alterations in human cancer. Nat. Biotechnol., 30, 413-421.

Cheng, D.T. et al. (2015) Memorial Sloan Kettering-Integrated Mutation Profiling of Actionable Cancer Targets (MSK-IMPACT): A Hybridization Capture-Based Next-Generation Sequencing Clinical Assay for Solid Tumor Molecular Oncology. J. Mol. Diagn., 17, 251-264.

Chen, L. et al. (2017) DNA damage is a pervasive cause of sequencing errors, directly confounding variant identification. Science, 355, 752-756.

Cibulskis, K. et al. (2013) Sensitive detection of somatic point mutations in impure and heterogeneous cancer samples. Nat. Biotechnol., 31, 213-219.

Garraway, L.A. (2013) Genomics-Driven Oncology: Framework for an Emerging Paradigm. J. Clin. Oncol., 31, 1806-1814.

Griffith, M. et al. (2017) CIViC is a community knowledgebase for expert crowdsourcing the clinical interpretation of variants in cancer. Nat. Genet., 49, 170-174.

Illumina (2017) Effects of Index Misassignment on Multiplexing and Downstream Analysis. URL: https://www.illumina.com/science/education/minimizing-index-hopping.html

Landau, D.A. et al. (2013) Evolution and impact of subclonal mutations in chronic lymphocytic leukemia. Cell, 152, 714-726.

Lee, J.-K. et al. (2016) Mechanisms and Consequences of Cancer Genome Instability: Lessons from Genome Sequencing Studies. Annu. Rev. Pathol., 11, 283-312.

McGranahan, N. et al. (2015) Clonal status of actionable driver events and the timing of mutational processes in cancer evolution. Sci. Transl. Med., 7, 283ra54.

Ryu, D. et al. (2016) Deciphering intratumor heterogeneity using cancer genome analysis. Hum. Genet., 135, 635-642.

Saunders, C.T. et al. (2012) Strelka: accurate somatic small-variant calling from sequenced tumor-normal sample pairs. Bioinformatics, 28, 1811-1817.

Schmitt, M.W. et al. (2016) The influence of subclonal resistance mutations on targeted cancer therapy. Nat. Rev. Clin. Oncol., 13, 335-347.

Sinha, R. et al. (2017) Index Switching Causes ‘Spreading-Of-Signal’ Among Multiplexed Samples In Illumina HiSeq 4000 DNA Sequencing. bioRxiv, 125724.

Zehir, A. et al. (2017) Mutational landscape of metastatic cancer revealed from prospective clinical sequencing of 10,000 patients. Nat. Med., 23, 703-713.

